# DYNLL1 mis-splicing is associated with replicative genome instability in SF3B1 mutant cells

**DOI:** 10.1101/2021.05.26.445839

**Authors:** Annie S. Tam, Shuhe Tsai, Emily Yun-Chia Chang, Veena Mathew, Alynn Shanks, T. Roderick Docking, Arun Kumar, Delphine G. Bernard, Aly Karsan, Peter C. Stirling

**Affiliations:** Terry Fox Laboratory, BC Cancer Agency. Vancouver, Canada; Department of Medical Genetics, University of British Columbia. Vancouver, Canada; Canada’s Michael Smith Genome Sciences Centre, BC Cancer, Vancouver, British Columbia, Canada; Université Brest, Inserm, EFS, UMR1078, GGB, F-29200 Brest, France; Department of Pathology and Laboratory Medicine, University of British Columbia, Vancouver, British Columbia, Canada

**Author notes:** Correspondence to P.C.S.

**Keywords:** SF3B1, DYNLL1, genome instability, R-loops, alternative splicing

## Abstract

Genome instability is a hallmark of cancer that arises through a panoply of mechanisms driven by oncogene and tumour-suppressor gene mutations. Oncogenic mutations in the core splicing factor SF3B1 have been linked to genome instability. Since SF3B1 mutations alter the selection of thousands of 3’ splice sites affecting genes across biological pathways, it is not entirely clear how they might drive genome instability. Here we confirm that while R-loop formation and associated replication stress may account for some of the SF3B1-mutant genome instability, a mechanism involving changes in gene expression also contributes. An SF3B1-H662Q mutant cell line mis-splices the 5’UTR of the DNA repair regulator DYNLL1, leading to higher DYNLL1 protein levels, mis-regulation of DNA repair pathway choice and PARP inhibitor sensitivity. Reduction of DYNLL1 protein in these cells restores genome stability. Together these data highlight how SF3B1 mutations can alter cancer hallmarks through subtle changes to the transcriptome.

## INTRODUCTION

*SF3B1* is the most frequently mutated splicing factor in cancer. Mutations in *SF3B1* occur in a broad range of cancers and blood disorders including myelodysplastic syndrome (MDS) (1, 2), chronic lymphocytic leukemia (CLL) (3, 4), acute myeloid leukemia (AML) (5), breast cancer (6) and uveal melanoma (7), amongst others (8). This implicates spliceosome dysfunction as a broad based driver of disease. The hotspot mutations identified in *SF3B1* are located in the HEAT repeat domain, and are heterozygous missense mutations, suggesting that these mutations likely confer an oncogenic gain of function phenotype rather than a loss of SF3B1 function (9). Many transcriptome analyses in different cancer types have characterized aberrant splicing events caused by *SF3B1* mutations (6, 7, 10–15), and although many overlapping events were observed across cancer types, the sets of aberrantly spliced junctions appear to cluster within each cancer type. These data seem to suggest that tissue specific splicing regulation may contribute to these aberrant splicing patterns. Functional consequences have in a few cases been assigned to SF3B1-mutant induced alternative splicing. For example, ABCB7 alternative splicing in SF3B1-mutant MDS has been proposed to deplete ABCB7 protein and contribute to the mitochondrial iron accumulation phenotypes in MDS with ring sideroblasts (16). Similarly, BRD9 alternative splicing in uveal melanoma leads to BRD9 depletion and loss of the tumor-suppressive functions of the non-canonical BAF complex (17). Thus, specific splicing changes clearly contribute to cancer hallmarks in SF3B1 mutant cancers. While these two important examples exist, the extent of functional splicing changes in SF3B1 mutant cancers is poorly understood overall.

Prior studies have linked SF3B1 mutations, and spliceosome disruption in general, to genome stability maintenance. For SF3B1, this could occur either through direct interactions with DNA damage response proteins like BRCA1 and BCLAF1 upon DNA damage (18), changes in expression of select genes like *ATM* and *TP53* which has been suggested to alter ATM/p53 transcriptional and apoptotic responses to DNA damaging agents(19), or global deregulation of splicing in factors that are important for cell cycle progression and DNA damage response (20). Another common mechanism of genome instability in cells with splicing defects is the accumulation of transcription coupled R-loops. These three-stranded R-loop structures consist of a DNA:RNA hybrid and exposed ssDNA on the non-template strand (21). R-loops contribute to genome instability by exposing ssDNA and by blocking replication forks, causing replication stress induced genome instability (22, 23). Previous work has suggested loss of splicing factors like ASF/SF2 (24), or treatment with splicing inhibitors induce aberrant R-loops (25, 26). In yeast, splicing has been shown to be an important factor in genome maintenance in this context as well (27, 28). Recently, two studies have demonstrated that cancer-associated mutations in splicing factors *U2AF1* and *SRSF2* elevated R-loops in leukemia cell lines, resulting in replication stress, activation of the ATR-CHK1 signaling pathway, and increased DNA damage (29, 30). Even more recently, two groups showed that CD34+ cells engineered to express SF3B1 mutants accumulated R-loops, DNA replication stress and sensitivity to ATR kinase inhibitors(31, 32). Understanding and possibly exploiting mechanisms of genome instability in SF3B1 mutant cancers is therefore of considerable ongoing interest.

In this study we focussed on an isogenic SF3B1 WT and mutant cell line pair to further probe genome stability mechanisms. We confirmed that R-loops are likely to accumulate in SF3B1 mutant cells. In addition, we observed induction of replication stress markers, decreased replication rates and overall replication dependence of genome instability in SF3B1 mutant cells. However, R-loops were not sufficient to explain all of the observed DNA damage. Another possibility is alternative splicing of transcripts encoding genome stability regulators and here we focus on DYNLL1 which is alternatively spliced in SF3B1 mutant cancers and our cell line models (6, 16). DYNLL1 encodes a small protein that homodimerizes and binds to dyneins, the BCL2 family protein BIM, and the DNA repair proteins 53BP1 and MRE11 (33–36). Due to its role in modulating DNA repair pathway choice there is growing interest in DYNLL1 loss as a chemoresistance mechanism (35–37). Here we associate DYNLL1 mis-splicing with DNA damage and instability in SF3B1 mutant cell line models, suggesting that alternative splicing promotes a gain of function and PARP inhibitor sensitization. These data highlight another potential mechanism of genome instability in SF3B1 mutant cancers.

## RESULTS AND DISCUSSION

### R-loops and DNA damage accumulate in SF3B1-mutant cell lines

To study the consequences of SF3B1 mutations, we screened two engineered leukemia cell lines K562 and NALM6 carrying SF3B1 hotspot mutations K700E and H662Q, respectively, to directly compare genome stability phenotypes with their isogenic wildtype controls. In our hands the NALM6 cells showed low levels of DNA damage in the control, while the *SF3B1*^*H662Q*^ mutant showed a clear accumulation of DNA breaks by the neutral comet assay (**Figure 1A**), and a significant increase in spontaneous γH2AX foci (**Figure 1B**). Single-stranded DNA damage was also evident in the alkaline comet assay (**Figure S1A**). Since the Boultwood lab recently established that *SF3B1*^*K700E*^ bearing cells accumulated R-loops, we next tested for DNA:RNA hybrids using the S9.6 monoclonal antibody in the NALM6 *SF3B1*^*H662Q*^ derivative. Immunofluorescence showed a significant increase in S9.6 staining in the *SF3B1*^*H662Q*^ mutants (**Figure 1C**). Given the potential role for R-loops in driving DNA damage, we next tested the effects of ectopic RNaseH1 expression, which should reduce R-loops, on the observed DNA damage phenotypes in NALM6 cells. GFP-RNaseH1 expression clearly reduced S9.6 signal in both the control and *SF3B1*^*H662Q*^ cells (**Figure 1C**), but did not significantly reduce the increased DNA breaks observed in the mutant (**Figure 1D**). While the DNA damage does seem to trend lower when RNaseH1 is overexpressed, these results suggest that R-loops may not be wholly responsible for the observed increases in DNA damage markers in *SF3B1*^*H662Q*^ cells. We thought that perhaps undamaged G1 cells were obscuring subtle differences but when we used EdU co-staining to separate G1 from S/G2 cells we still found a significant increase in DNA damage in *SF3B1*^*H662Q*^ cells that was not completely suppressed by RNAseH1 expression (**Figure 1E**). Interestingly, this analysis showed no difference in γH2AX foci for control and *SF3B1*^*H662Q*^ cells in G1 phase, suggesting a DNA replication-based mechanism of damage that we decided to explore further.

**Figure 1.**
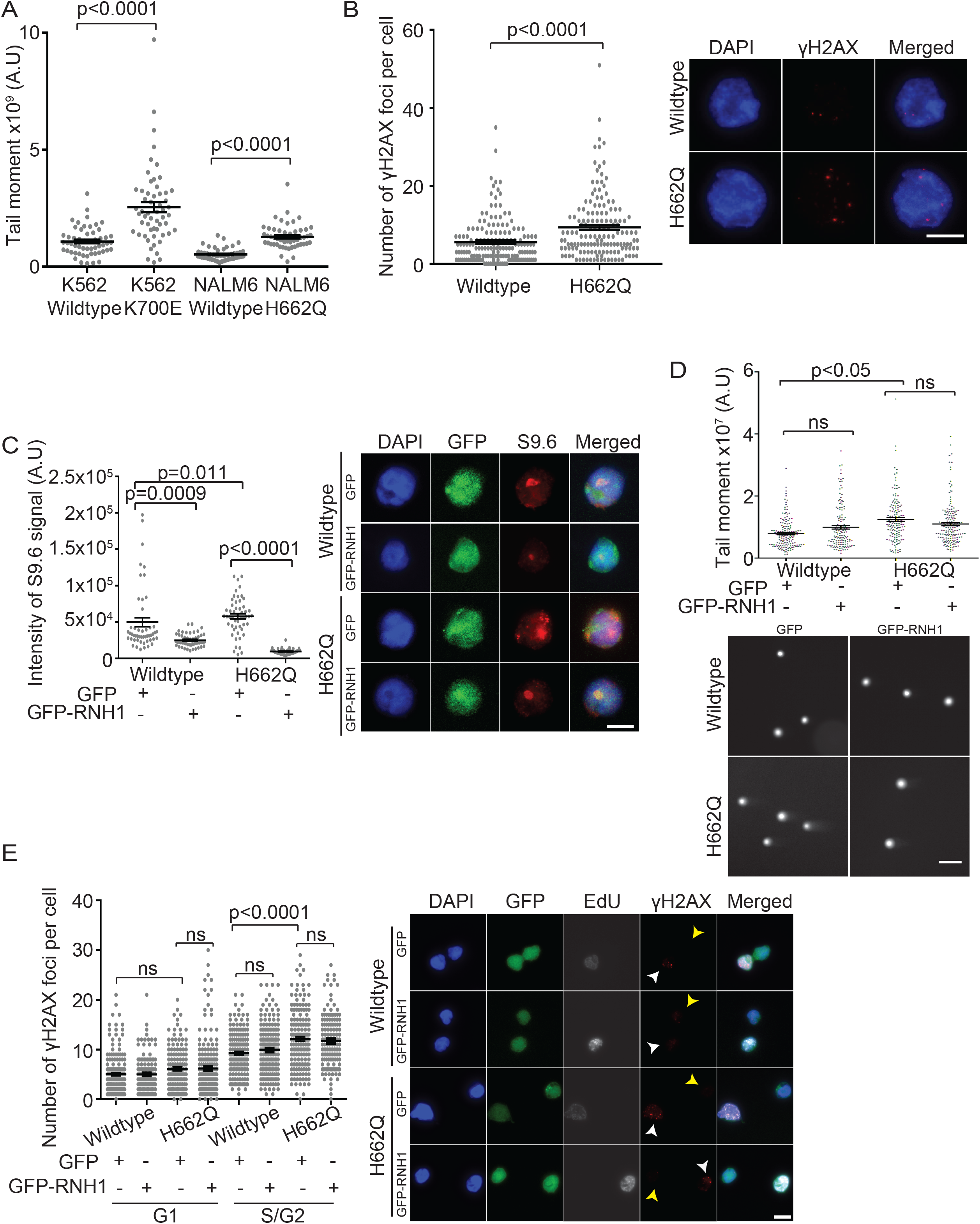
DNA damage and R-loops in SF3B1 mutant cell lines. (A) Neutral comet assay for two isogenic cell line pairs with the indicated genotypes. >50 cells scored per replicate. (B) Quantification of γH2AX foci in the NALM6 control and *SF3B1*^*H662Q*^ lines. >100 nuclei scored per replicate. (C) Intensity of S9.6 antibody staining in the NALM6 cell line pair with either control vector (GFP) or one expressing GFP-RNaseH1 (GFP-RNH1). >100 GFP positive nuclei scored per replicate. (D) Neutral comet assay of cells transfected with GFP control vector or GFP-RNaseH1. >50 cells scored per replicate. (E) Quantification of γH2AX foci in EdU positive versus negative cells with or without ectopic GFP-RNaseH1 expression. >100 GFP positive nuclei scored per replicate. Yellow arrows indicate EdU negative (G1 phase) cells; White arrows indicate EdU positive (S/G2 phase) cells. (A-E) Mean values with S.E.M. error bars are shown, *n = 3*. (B-E) Representative image scale bars = 5 μm. Student’s t-test (A-B) and ANOVA with Tukey’s post-hoc test (C-E) were used to calculate statistical significance.

### DNA replication stress in SF3B1^H662Q^ cells

To probe the potential role of DNA replication in inducing DNA damage in the *SF3B1*^*H662Q*^ cells we first used western blot to measure the output of ATR signaling in the cells. Phosphorylation of the apical replication stress sensor ATR and its effector kinase Chk1 were consistently higher in the *SF3B1*^*H662Q*^ cells (**Figure 2A**). Accordingly, phosphorylation of the single-stranded DNA binding protein RPA32 was also increased, while ATM and Chk2 phosphorylation were not dramatically affected (**Figure 2A**). *SF3B1*^*K700E*^ K562 cells also showed higher levels of RPA32-ser33P indicative of replication stress (**Figure S1B**). Using RPA32-serine 33 phosphorylation as a marker of DNA replication stress we performed immunofluorescence in concert with EdU staining and RNaseH1 overexpression. These data again showed that RPA32-Ser33P foci increased in both EdU- and EdU+ *SF3B1*^*H662Q*^ cells (**Figure 2B**). Importantly, ectopic GFP-RNaseH1 expression significantly reduced the phosphorylated RPA32 foci in both conditions. Previous work has suggested that RPA binds directly to single-stranded DNA exposed by R-loop structures (38, 39). Moreover, it is possible that some EdU-cells are in fact arrested S-phase cells that do not incorporate sufficient EdU during the time course of the experiment. Thus, RNaseH1 might reduce the phosphorylated RPA32 foci by resolving R-loops associated with RPA, and by mitigating replication stress associated with transcription-replication conflicts. Regardless, these data show that some replication stress phenotypes are likely to be R-loop dependent, even though the persistence of DNA breaks (**Figure 1**) is not wholly sensitive to RNaseH. Consistent with the observed markers of replication stress, we directly observed reduced replication speed in *SF3B1*^*H662Q*^ cells by DNA combing (**Figure 2C**). We also observed a small but significant increase in the proportion of S-phase *SF3B1*^*H662Q*^ cells by flow cytometry, suggesting that the defects are not triggering a complete cell cycle arrest (**Figure S1C**). However, failure to complete DNA replication can lead to under-replicated and concatenated chromosomes that are marked by anaphase bridges when cells attempt to undergo mitosis. Consistent with SF3B1 mutations inducing chromosome instability in this NALM6 cell model, we saw a significant increase in BLM helicase-marked anaphase bridges in DAPI-stained anaphase cells (**Figure 2D**). Finally, to confirm that DNA replication is the major driver of DNA breaks in the *SF3B1*^*H662Q*^ cells, we inhibited the Cdc7 kinase with PHA-767491 (Cdc7i), a potent inhibitor of CDC7(40) that ablates MCM2 phosphorylation (**Figure S1D**) and subsequent replication initiation. Cdc7 inhibition completely suppressed DNA breaks as measured by the neutral comet assay (**Figure 2E**) and blocked mutant induced RPA32 phosphorylation by western blot (**Figure S1E**). Our results thus far confirm that SF3B1 mutant cells accumulate DNA damage and exhibit genome instability. This damage is dependent on DNA replication and is associated with a DNA replication stress response. The role of R-loops in these phenotypes is not completely penetrant, although we do see effects consistent with reports linking stabilized R-loops to splicing factor mutations (29, 31, 32). While R-loops may be a consequence of splicing disruption, since RNaseH1 could not completely suppress DNA strand breaks in the SF3B1 mutant model, we sought additional causes of genome instability.

**Figure 2.**
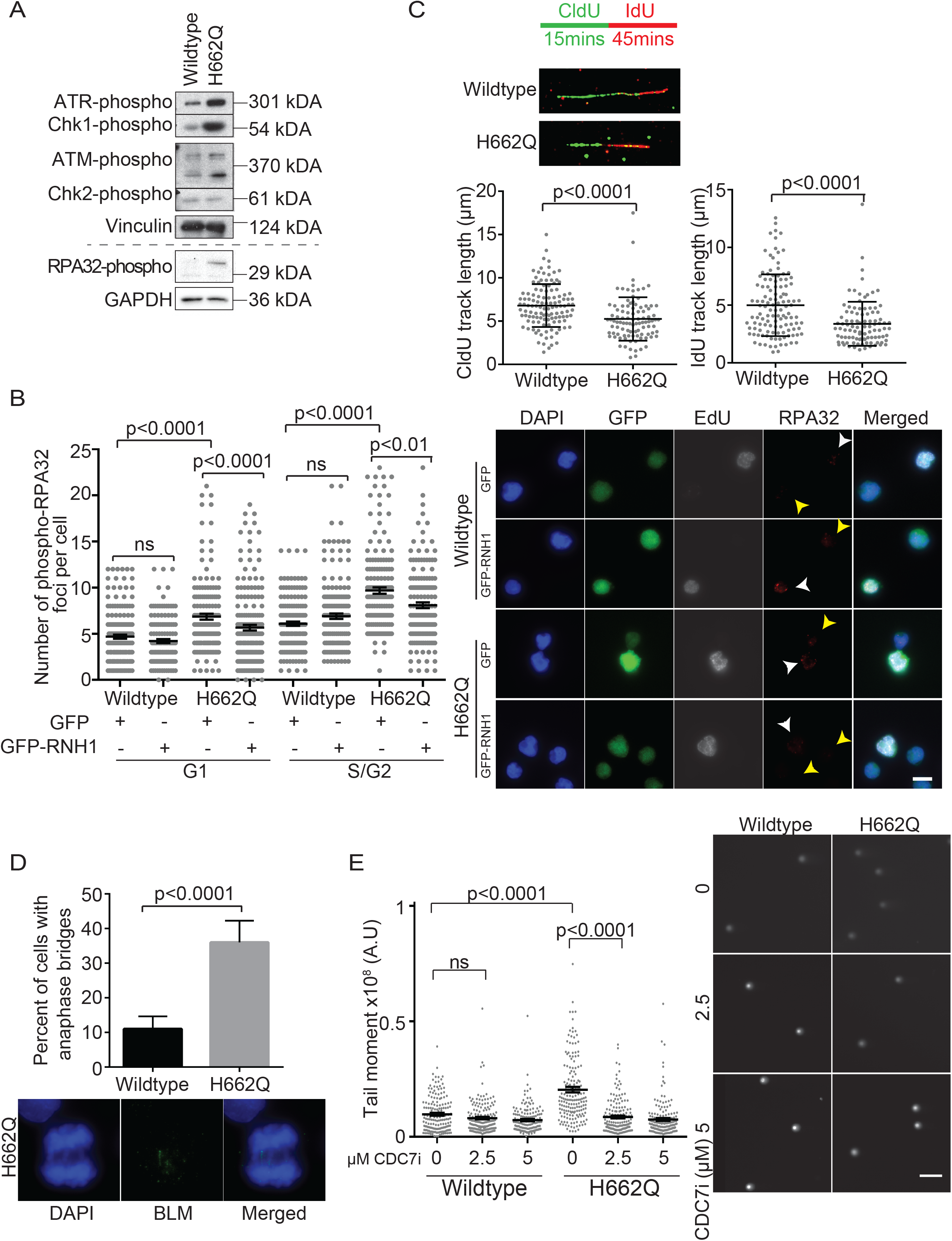
DNA replication is the driver of DNA damage in SF3B1^H662Q^ cells. (A) Representative western blot analysis of the indicated proteins in cycling NALM6 control and *SF3B1*^*H662Q*^ cells; *n = 3*. (B) Immunofluorescence of phosphorylated RPA32-Ser33 in the indicated EdU+ or EdU-cells with control (GFP) or with ectopic GFP-RNaseH1 expression. >100 GFP positive nuclei scored per replicate. Scale bar = 5 μm. Yellow arrows indicate EdU negative (G1 phase) cells; White arrows indicate EdU positive (S/G2 phase) cells. ANOVA with Tukey’s post-hoc test was used to determine statistical significance. Mean values with S.E.M. error bars are shown, *n = 3*. (C) Analysis of DNA replication fork speed in NALM6 and *SF3B1*^*H662Q*^ cells by DNA fiber combing. Replication forks were labelled with CldU (green) and IdU (red). Unpaired t-test was used to determine statistical significance. Mean values with S.D. are shown, *n = 2*. 120 and 98 DNA fibers were measured for WT and mutant, respectively. (D) Quantification of BLM helicase-positive anaphase bridges in the indicated cells. Fisher’s exact test was used to determine statistical significance. Mean values with S.E.M. error bars are shown, *n =4*. 175 anaphase cells were scored per cell line. (E) Neutral comet assay of NALM6 or *SF3B1*^*H662Q*^ cells in the presence of Cdc7 inhibitor. >50 cells scored per replicate. ANOVA with Tukey’s post-hoc test was used to determine statistical significance. Mean values with S.E.M. error bars are shown, *n = 3*. Scale bar = 5 μm.

### DYNLL1 mis-splicing in SF3B1^H662Q^ cells is associated with higher DYNLL1 protein

Another potential mechanism of genome instability is that aberrant splicing products deplete proteins essential for genome maintenance. We have shown that this is the case for α-tubulin production using a yeast model (28). There is also precedent for single gene mis-splicing effects in the case of SF3B1 mutated cancers. Aberrant splicing of the iron transporter *ABCB7* has been suggested to contribute to the buildup of erythroid precursors with abnormal mitochondrial iron accumulation (ring sideroblasts) in SF3B1-mutated MDS (16, 41). Despite large changes in the transcriptome from individual published datasets, analysis of common splicing changes from several RNA sequencing studies annotates universal splicing changes in the genes *ABCB7, ABCC5, DYNLL1, TMEM14C* and *SEPT6* (7, 12, 15, 42). Among these only DYNLL1 has a well described role in DNA damage repair (35, 36). In addition to binding to cytoplasmic dynein and the apoptotic regulator Bim, DYNLL1 binds to the MRE11 nuclease and negatively regulates its nuclease activity while also promoting the activity of 53BP1, and therefore plays a role in regulating DNA double strand break repair pathway choice (35, 36). SF3B1 mutations have been associated with inclusion of an additional 14bp in the 5’UTR of the DYNLL1 transcript (**Figure 3A**), and this long variant isoform is found with high percent spliced-in in cancer transcriptomes (9- or 17-fold increases in usage; (11, 12)). This change to a longer isoform is clearly evident in end-point RT-PCR reactions derived from *SF3B1*^*H662Q*^ mutant, but not WT NALM6 cell lines (**Figure 3B**). The same result is seen in *SF3B1*^*K700E*^ K562 cells (**Figure S2A**). Importantly, quantitative PCR confirms a significant splicing-in of the extended 14bp 5’UTR, but no overall changes in DYNLL1 total transcript levels as measured across three exons (**Figure 3C**). Moreover, retrospective analysis of RefSeq transcript usage in RNA-seq data from an MDS/AML cohort (n=173) (43) shows that SF3B1 mutant cases express significantly higher levels of the long transcript than average cases with wildtype SF3B1 (**Figure 3D**). Thus, our data confirms that multiple SF3B1 mutations are associated with alternative splicing of DYNLL1 and DNA damage phenotypes.

**Figure 3.**
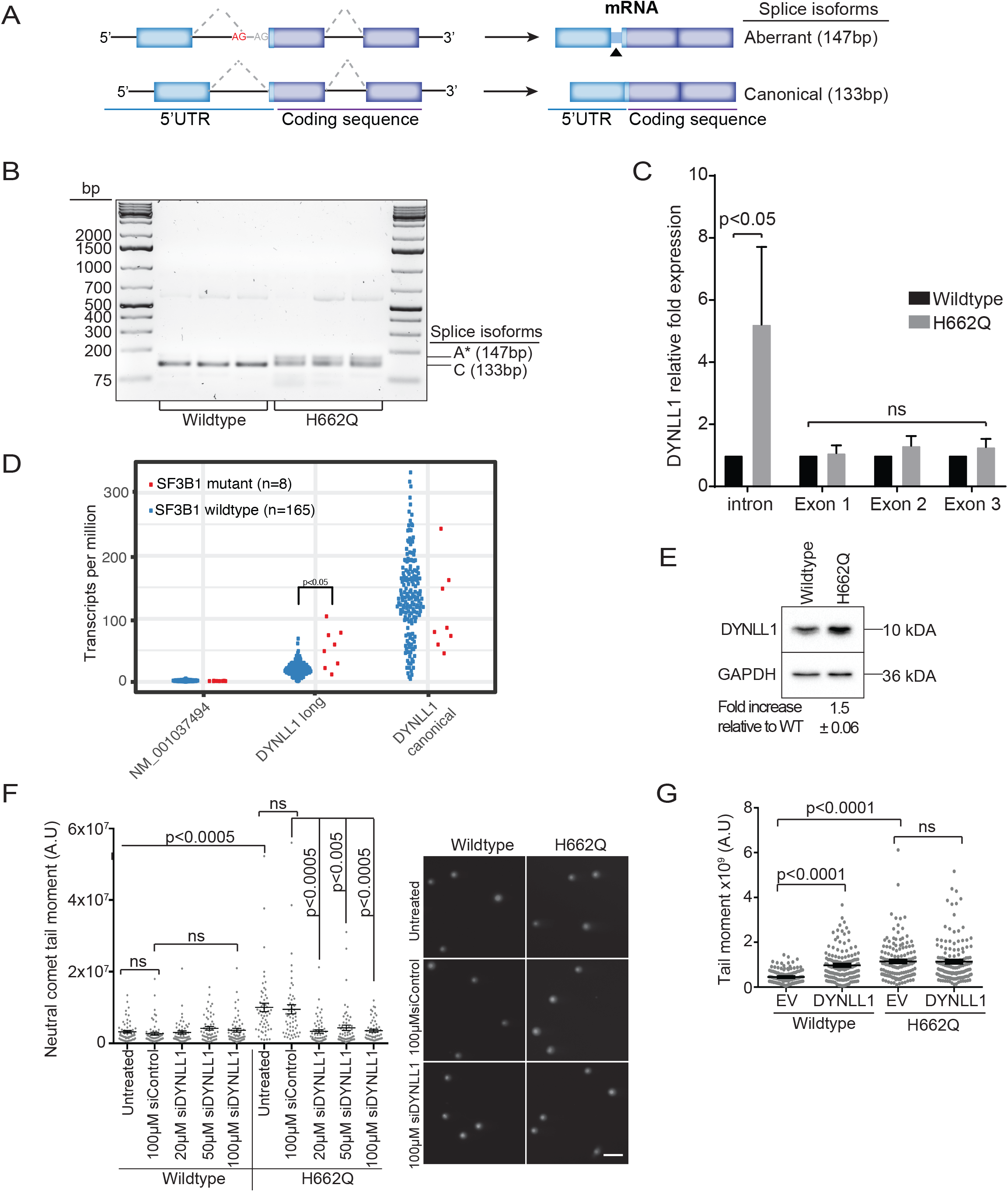
Alternative splicing and overexpression of DYNLL1 *SF3B1* mutant cells. (A) Schematic of short and long isoforms of DYNLL1 showing the 14-bp change in the 5’UTR. (B) Endpoint RT-PCR analysis of DYNLL1 transcripts in the indicated cell lines. Technical triplicates are shown for each cell line. C = canonical transcript, A* = aberrant transcript. (C) Relative fold expression of DYNLL1 transcript normalized to ACTB measured by four sets of qPCR primers. ‘Intron’ only captures the incorporation of the 14bp DYNLL1 long segment, while the Exon 1, 2, and 3 primers measure total transcript. Three biological replicates; Mean values with S.E.M. error bars; p-values are derived from ANOVA followed by a Holm-Sidak post-hoc test. (D) Analysis of DYNLL1 RefSeq transcript usage in 173 RNA sequencing datasets for MDS or AML cases with known SF3B1 mutation status(43). Shown are a non-expressed variant NM_001037494, the canonical transcript NM_00376, and the long 5’UTR variant NM_001037495 (DYNLL1 long). Red points are SF3B1-mutant cases. The indicated p-value compares transcript expression between mutant and wildtype SF3B1 cases using a two-sided *t* test. (E) Representative western blot of DYNLL1 protein, quantification of triplicate experiments is shown below. (F) Neutral comet assay after si-DYNLL1 or control siRNA knockdown treatment in the indicated cells. Mean values with S.E.M. error bars are shown, *n = 3*. Scale bar = 5 μm. (G) Neutral comet assay of cells with an empty control vector (EV) or a DYNLL1 overexpression plasmid. >50 cells scored per replicate. Mean values with S.E.M. error bars are shown, *n = 3*. For E and F, ANOVA with Tukey’s post-hoc test was used to determine statistical significance. Ns = not significant.

Since the extended 5’UTR should not change the coding sequence of DYNLL1 we wondered how this could affect DYNLL1 protein levels. Western blotting for DYNLL1 protein revealed that either *SF3B1*^*H662Q*^ NALM6 mutant cells, or *SF3B1*^*K700E*^ K562 mutant cells had higher levels of DYNLL1 protein (**Figure 3E** and **Figure S2B**). To directly assess whether high levels of DYNLL1 protein could be responsible for the observed DNA damage we reduced DYNLL1 levels with a low dose of DYNLL1 siRNA and retested cells for damage using the neutral comet assay (**Figure 3F**). DYNLL1 siRNA treatment suppressed the additional DNA breaks in the *SF3B1*^*H662Q*^ cells. Together these data show that a DYNLL1 long-5’UTR isoform is expressed in SF3B1 mutant cells, and that this correlates with overexpression of DYNLL1 protein. Reducing DYNLL1 protein with siRNA reverses DNA damage, suggesting that high DYNLL1 protein may be a cause of DNA damage. To confirm this we transiently transfected a cDNA to overexpress DYNLL1 in NALM6 cells. The DYNLL1 cDNA increased DNA breaks as measured by comet assay in WT NALM6 cells but not in the SF3B1 mutant line (**Figure 3G**). We also took advantage of a recently reported DOX-inducible *SF3B1*^*K700E*^ model derived from K562 cells (44). Induction of *SF3B1*^*K700E*^ but not WT SF3B1 with DOX for 48 hours also led to accumulation of the long DYNLL1 transcript by RT-PCR (**Figure S3A** and **S3B**). Importantly at the same time we saw induction of RPA32-ser33P foci by immunofluorescence only in the *SF3B1*^*K700E*^ but not in SF3B1 WT induced cells (**Figure S3C**). These analyses further support a correlation between SF3B1-mutant alternative splicing of DYNLL1 and DNA damage. Overall, SF3B1 alternative splicing of DYNLL1 may modulate its protein levels or availability, and our data suggest that the genome is highly sensitive to DYNLL1 protein abundance.

### Shifts in DNA repair pathway choice and Olaparib sensitivity in SF3B1^H662Q^ cells

DYNLL1 has been shown to have at least two roles in regulating the balance between NHEJ and HR. First, it was found that DYNLL1 specifically binds to 53BP1 (45), and this interaction plays an essential role in promoting NHEJ, because depletion of DYNLL1 reduced the efficiency of 53BP1-mediated NHEJ (35). Second, DYNLL1 directly interacts with MRE11 to limit DNA end resection, leading to reduced HR pathway choice (36). To determine if SF3B1 mutations alter the balance of these pathways, we performed immunofluorescence experiments co-staining EdU treated cells with 53BP1 antibody as a marker for NHEJ and RAD51 antibody to mark HR. The results show that *SF3B1*^*H662Q*^ mutant cells had a higher number of 53BP1 foci at G1 and S/G2 phases (**Figure 4A**), and a lower number of RAD51 foci at S/G2 phase relative to wildtype (**Figure 4B**). Since HR is generally active in S and G2 phases of the cell cycle when the sister chromatid is available as a repair template, the immunofluorescence data showing that RAD51 foci are predominantly observed in wildtype S/G2 phases is consistent with RAD51 as a marker for HR, and suggests that the mutant cells have a defect in normal repair pathway choice. Consistent with a role in modulating DSB repair choice, loss of DYNLL1 has been identified as a PARP-inhibitor (PARPi) resistance mechanism in HR-deficient cancer cell lines, and low DYNLL1 expression may influence genome stability in patient samples(35, 36). If DYNLL1 loss leads to PARPi resistance then we reasoned that overexpression of DYNLL1 should sensitize cells to PARPi. Accordingly, the *SF3B1*^*H662Q*^ cells were significantly more sensitive to olaparib than the parental WT cells (**Figure 4C**). Olaparib was also more toxic to *SF3B1*^*K700E*^ K562 cells compared to WT, although the effects were not as strong (**Figure S2C**). Since NALM6 cells are HR-proficient, this raises the possibility that SF3B1 mutations alone, or high expression of the DYNLL1-long transcript might provide a rationale for targeting tumours with PARP inhibitors (**Figure 4D**). Consistent with this prospect, a phase I clinical trial of olaparib in relapsed blood cancers has suggested better responses in patients with SF3B1 or ATM mutations albeit in a small cohort (46).

**Figure 4.**
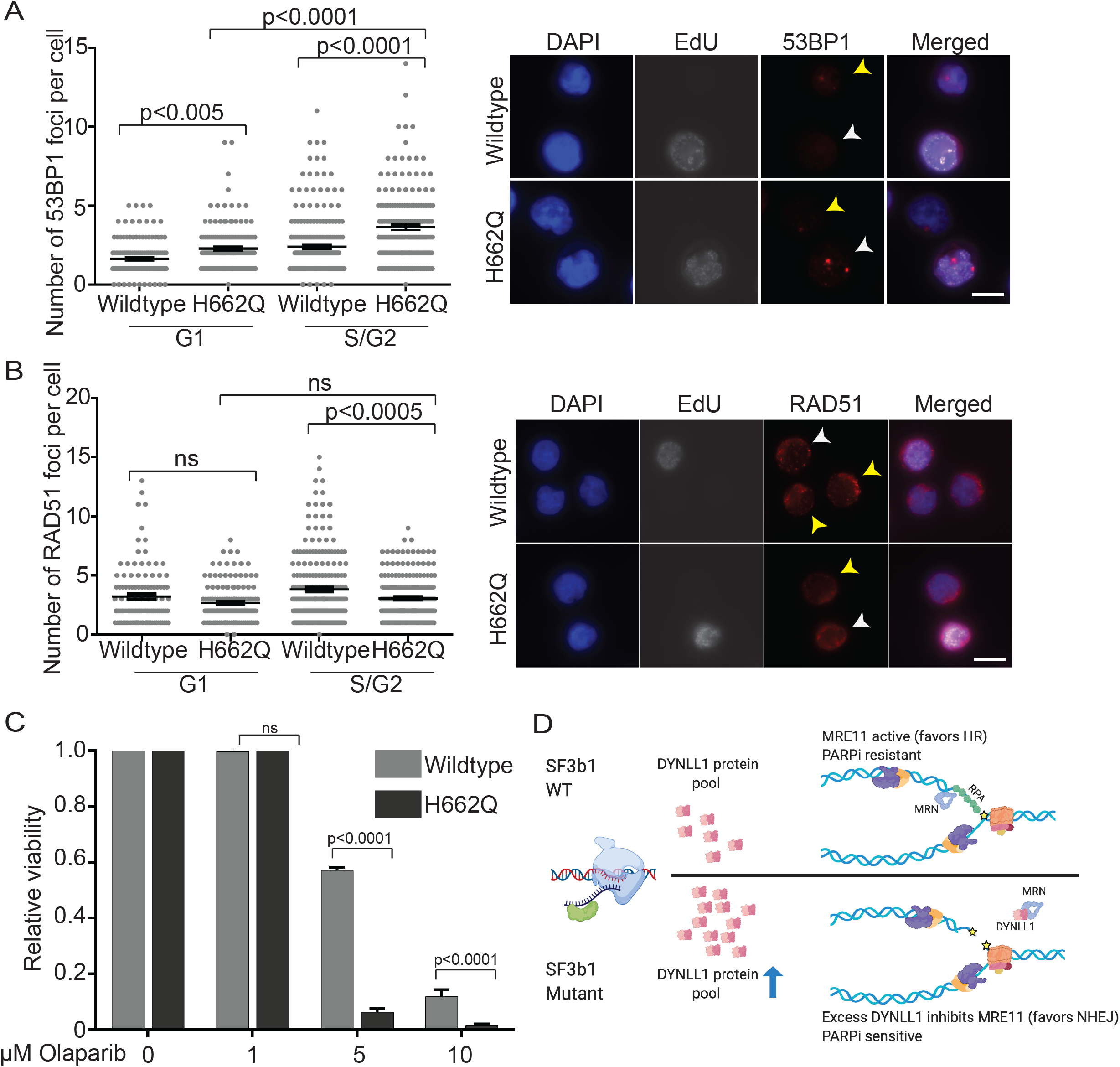
Alterations in DNA repair foci and olaparib sensitivity in *SF3B1*^*H662Q*^ cells. Immunofluorescence for (A) 53BP1 foci and (B) RAD51 foci after EdU incorporation. Mean values with S.E.M. error bars are shown, *n = 3*. >100 nuclei were scored per replicate. Scale bar = 5 μm. Yellow arrows indicate EdU negative (G1 phase) cells; White arrows indicate EdU positive (S/G2 phase) cells. (C) Relative cell viability of the indicated cell lines following olaparib treatment at the indicated doses. Mean values with S.E.M. error bars are shown, *n = 3*. For A-D, ANOVA with Tukey’s post-hoc test was used to determine statistical significance. (D) Model of SF3B1 mutant driven genome instability through DYNLL1. In SF3B1 WT cells (top) DYNLL1 protein pools are normal and cells facing replication blocks have normal MRE11 regulation, which favours HR repair in S/G2 cells and permits resistance to PARP trapping inhibitors that increase requirements for HR repair. In SF3B1 mutant cells (bottom), alternative splicing is associated with increased DYNLL1 protein pools. In this scenario cells facing replication challenges have abnormal MRE11 regulation due to excess DYNLL1 which limits fork restart and repair by HR mechanisms. This reduced HR and overactive NHEJ (53BP1 foci in A) sensitize SF3B1 mutant cells to PARP inhibition. E was created with Biorender.com

### Perspective

A panoply of factors with diverse cellular functions directly or indirectly regulate genome stability. Changes in RNA splicing have emerged as potential dual regulators of genome stability, since RNA can be directly involved in DNA damage and repair, and because of changes to gene expression due to mis-splicing of important transcripts (21, 22). Here we explore the genome instability phenotypes associated with one of the most common splicing alterations in cancer, hotspot mutations in SF3B1. Using isogenic cell line pairs we confirm that SF3B1 mutations H662Q and K700E are associated with increased DNA replication stress and R-loop accumulation. However, alternative splicing of the DNA repair regulator DYNLL1 also accumulates in SF3B1 mutant cells and our data suggests that this is associated with shifts in repair pathway choice and DYNLL1-dependent DNA damage. We find that DYNLL1 overexpression is sufficient to induce DNA damage and that the use of the DYNLL1-long transcript is associated with small but significant increases in DYNLL1 protein. Whether small changes in DYNLL1 protein are really enough to shift the DNA repair state in tumours is unknown. While the extended 5’UTR in the long DYNLL1 transcript does not produce predicted coding changes, it could affect mRNA localization, stability or translation in ways that alter DYNLL1 regulation at the RNA level. Given the strong association of DYNLL1 mis-splicing with SF3B1 mutations and the importance of DYNLL1 for genome maintenance and other cancer hallmarks, further work on the mechanism of dysregulation is warranted. Indeed, what other changes to the spliceosome cause DYNLL1 alternative-splicing is unknown. Recent work shows that loss of the splicing factor SUGP1 largely mimics the effects of SF3B1 hotspot mutations, suggesting an alternate mechanism that could affect DYNLL1 (47, 48). Whether other common splicing mutations in *U2AF1, SRSF2, ZRSR2*, or other factors influence DYNLL1 is unknown.

Previous work has found that cancer-associated mutations of SF3B1 and other splicing factors cause R-loop accumulation and associated genome instability (29, 31). Presumably these R-loops occur in cis to failed or slowed co-transcriptional splicing reactions, although the SF3B1 mutant containing spliceosomes must be largely functional to support cell viability. While unscheduled R-loop formation is certainly a driver of replication stress in some systems, our data support a mechanism in which changes to the transcriptome can also influence genome stability in parallel with R-loops. This must be considered as researchers discover an expanding set of transcriptional regulators, chromatin modifiers, and RNA processing factors that all seem to regulate R-loop associated DNA damage (21). Interestingly, some SF3B1 mutant cancers like MDS have a relatively low mutation burden suggesting that potential replication stress is generally managed without accumulating a lot of mutations. This is reminiscent of work on Ewing’s sarcoma which also showed high levels of R-loops, replication stress and PARP inhibitor sensitivity, in a tumour with very low levels of somatic mutations (49).

DYNLL1 through its effects on MRE11 sensitizes cells to PARP inhibitors by favouring non-homologous end joining (35). Much of this activity has been described in the context of DNA double strand break repair, but our data suggests that increased DNA damage due to functions for DYNLL1 during DNA replication may be important for understanding its effects on genome maintenance. In this light, MRE11 clearly plays both nucleolytic and structural roles at DNA replication forks that are important for genome maintenance. Therefore it is plausible that higher levels of nuclear DYNLL1 protein might influence key functions of MRE11 at forks. Learning how DYNLL1 is regulated and in-turn regulates MRE11 at replication forks is an important area for future research.

## ACKNOWLEDGEMENTS

This work was supported by the Canadian Institutes of Health Research (CIHR MOP136982). A.S.T. was supported by an Elizabeth C. Watters research fellowship. P.C.S is a CIHR New Investigator and a Michael Smith Foundation for Health Research Scholar. Work in the laboratory of AK is supported by grants from Genome British Columbia (X01CBP; #121AML) and the BC Cancer Foundation through the Leukemia and Myeloma Program (LaMP). AK is supported by the John Auston BC Cancer Foundation Clinician-Scientist Award.

## METHODS

### Cell lines, siRNA, plasmids, and cell culture

NALM6 wildtype and *SF3B1-H662Q* cells were purchased from Horizon Discovery and grown in RPMI-1640 medium (STEMCELL Technologies) supplemented with 10% fetal bovine serum (Life Technologies) in 5% CO_2_ at 37°C. Transfections were performed using the Neon™ Transfection System (Invitrogen) following manufacturer instructions. Briefly, 80-90% confluent cells were harvested and washed twice with PBS, then 2×10^5^ cells were transfected with a 10μL Neon Tip via electroporation using the following optimized settings: 1350 V, pulse length 10ms, total of 4 pulses. Cells were harvested 48 hours after transfection. For experiments with overexpression of GFP or nuclear-targeting GFP-RNaseH1 (gift from R. Crouch), 1 μg of plasmid per reaction was used. For RNA interference, siRNA concentrations of 20, 50, and 100 μM si-Control or si-DYNLL1 were used (siGENOME-SMARTpool siRNAs from Dharmacon). The DYNLL1 overexpression cDNA clone expressed from a pcDNA3 vector obtained from Addgene (Plasmid #24265 (50)) was compared to an empty pcDNA3 control. The DOX-inducible K562 cell line models were cultured as above and then treated with 2μg/mL of DOX for 48 hours before being subject to RT-PCR, western blot or immunofluorescence as described below.

### Western blot analysis

Whole-cell lysates were prepared using 1 x RIPA buffer containing protease inhibitor (Sigma) and phosphatase inhibitor PhosSTOP™ (Roche Applied Science) cocktail tablets. 10X RIPA buffer: 0.5M Tris-HCl, pH 7.4, 1.5M NaCl, 2.5% deoxycholic acid, 10% NP-40, 10mM EDTA. Protein concentrations were assessed with Bio-Rad Protein assay (Bio-Rad). Lysates were separated on a 15%, 10%, or 8%/15% gradient SDS-PAGE gel, transferred to 0.45μm PVDF membranes (Millipore), blocked with 5% skim milk in TBS containing 0.1% Tween-20 (TBS-T), and probed with the following antibodies: γH2AX (1:1000, abcam cat#ab81299), p-ATR (Ser428) (1:1000, Cell Signaling cat#2853), p-CHK1 (Ser345) (1:1000, Cell Signaling cat#133D3), p-ATM (1:500, Santa Cruz cat#sc-47739), p-CHK2 (Th468) (1:1000, Cell Signaling cat#C13C1), GAPDH (1:3000, ThermoFisher Scientific cat# MA5-15738), vinculin (1:2000, Santa Cruz cat#sc-73614), p-RPA32 (1:1000, Bethyl cat# A300-246A), RAD51 (1:500, Santa Cruz sc-398587), CDC7 (1:500, Santa Cruz, sc-56274), p-MCM2 (1:2000, Abcam, cat# ab70371), MCM2 total (1:1000, Abcam cat# ab108935), DYNLL1 (1:5000, Abcam, cat# ab51603). Secondary antibodies were conjugated to horseradish peroxidase (HRP) and peroxidase activity was visualized using SuperSignal™ West Pico PLUS Chemiluminescent Substrate (Thermo Scientific cat# 34577). ImageJ software was used to quantify protein bands(51).

### Immunofluorescence and EdU staining for imaging

Suspension cells were harvested, washed twice with PBS and incubated on poly-L-lysine coated coverslips for 15 mins at room temperature before fixation. To make coverslips, 0.1% poly-L-lysine solution (Sigma-Aldrich cat# P8920) was added to coverslips for 20 mins, removed, and slides were dried at 37°C for 15 mins. For S9.6 staining, cells were fixed with ice-cold methanol for 15 mins and permeabilized with ice-cold acetone for 1 min. After PBS wash, cells were blocked in 3% bovine serum albumin (BSA), 0.1% Tween-20 in 4X saline sodium citrate buffer (SSC) for 30 mins at room temperature in a humid chamber. Cells were incubated with primary antibody DNA-RNA Hybrid [S9.6] (1:500, Kerafast cat#ENH001) overnight at 4°C. Next day, cells were washed three times with PBS, and stained with secondary Alexa Fluor® 568 goat anti-Mouse IgG (1:1000, Invitrogen cat#A-11004) for 1 hr at room temperature, washed twice and PBS and one time with PBS+0.1% Tween-20 (PBS-T), and stained with DAPI for 5 mins. Cells were imaged on a LeicaDMI8 microscope at 100X and ImageJ was used for processing and quantifying nuclear S9.6 intensity in images. For experiments with GFP overexpression, only GFP-positive cells were quantified. For γH2AX (1:1000), MCM2 (1:1000), RPA32 (1:1000), 53BP1 (1:1000), and RAD51 (1:500), immunostaining was performed as above, except for the following: fixation with 4% paraformaldehyde for 15 mins and permeabilization with 0.2% Triton X-100 for 5 mins; secondary antibody was rabbit or mouse Alexa-Fluoro-488 or 568-conjugated antibody (1:1000, Invitrogen). For EdU incorporation, Click-iT™ EdU Imaging Kit with Alexa Fluor™ 647 Azides (Invitrogen cat# C10086) was used following manufacturer instructions. Cells were labeled with 10 μM EdU for 15 mins before fixation. After fixation and permeabilization, Click-iT® reaction cocktail with Alexa Fluor™ 647 was added to slides for 30 mins, wash twice with 3% BSA in PBS, and blocked with 3% bovine serum albumin (BSA), 0.1% Tween-20 in 4X saline sodium citrate buffer (SSC) for 30 mins at room temperature in a humid chamber. Antibody staining proceeded as described above.

### Neutral and alkaline comet assays

The neutral and alkaline comet assays were performed using the CometAssay Reagent Kit for Single Cell Gel Electrophoresis Assay (Trevigen cat#4250-050-K) following manufacturer instructions. Cells at a concentration of 1.5 × 10^5^ cells/mL were combined with low melt agarose (LMAgarose) at 37 °C at a ratio of 1:10 (cell:LMAgarose) and spread onto CometSlide. After gelling in 4 °C in the dark, the slides were then immersed in Lysis Solution overnight at 4°C. For the neutral comet assay, slides were removed from Lysis Solution and immersed in 4°C Neutral Electrophoresis Buffer for 30 min. Electrophoresis was then performed at 4°C at 21 Volts for 40 min in Neutral Electrophoresis Buffer. Slides were immersed in DNA Precipitation Solution for 30 min followed by 70% ethanol for 30 min at room temperature, and dried at 37°C for 15 min. Slides were then stained with 16 μg/mL propidium iodide (Sigma-Aldrich cat#287075) for 30 mins, gently washed with water, and allowed to dry completely at 37°C for 15 min. Slides were imaged on LeicaDMI8 microscope at 10X. Comet tail moments (tail length x fraction of total DNA in the tail) were obtained using an ImageJ plugin as previously described (52), and at least 50 cells per sample were analyzed for each independent replicate. For the alkaline comet assay, the protocol is the same except for the following: after lysis, slides were immersed in Alkaline Unwinding Solution for 20 mins at room temperature, electrophoresis was done in Alkaline Electrophoresis Solution, after which the slides were immersed twice in dH2O for 5 mins each, followed by 70% ethanol for 5 mins.

### Cell viability measurements

Cell numbers and viability were measured using the Advanced Image Cytometer NucleoCounter® NC-250™ (ChemoMetec). 20 µL of cell suspension was incubated with 1 µL of Solution 18 (ChemoMetec cat#910-3018) in Eppendorf tubes, and 10 µL of cell:dye mixture was added to NC-Slide A8™ chamber slides (ChemoMetec cat# 942-0003) for analysis. Solution 18 contains acridine orange (AO) which is used to counterstain living and dead cells, and DAPI, which is used to stain dead cells. For cell viability analysis, 500 µL of cell cultures were incubated for 72 hours, and measurements were taken every 24 hours. For cell viability and doubling capacity post-drug treatment, cells were incubated at the respective drug concentrations for 72 hours, except in the case of olaparib and mirin treatments where cells were initially treated for 48 hours, then media and drugs were replenished for another 72 hours. For viability measurements post si-DYNLL1, 500 µL of cell cultures were treated with 0, 20, 50, or 100 µM si-Control or si-DYNLL1 for 48 hours.

### Cell cycle analysis

Cell cycle progression of NALM6 cells was assessed using Click-iT™ EdU Alexa Fluor™ 488 Flow Cytometry Assay Kit (Invitrogen cat # C10425), following manufacturer instructions. Cells were incubated with 10 µM EdU for 45 mins, harvested, and washed twice with PBS. Cells were washed once with 1% BSA in PBS, fixed with 100 µL of Click-iT® fixative for 15 mins at room temperature, washed with 1% BSA in PBS, and permeabilized with 100 μL of 1X Click-iT® saponin-based permeabilization and wash reagent for 15 mins at room temperature. Click-iT® reaction cocktail (PBS, CuSO4, Alexafluor-488-azide, 1x Reaction Buffer Additive) was added to cells for 30 mins at room temperature protected from light. Lastly, cells were washed with 1X Click-iT® saponin-based permeabilization and wash reagent, and stained with 16 μg/mL propidium iodide (Sigma-Aldrich cat#287075) with 0.25 mg/mL Ribonuclease A for 30 mins at room temperature. Cells were washed and resuspended in PBS and analyzed with a BD LSRFortessa™ flow cytometer using CellQuestPro software. Cell cycle plots were generated using FlowJo Version 9.3.2.

### Drug treatments

NALM6 cells were plated at a concentration of 6 × 10^5^ cells/mL, 500 μL per well in a 24-well plate. Cells were cultured with the following concentrations of drugs: CDC7 inhibitor PHA-767491 (Selleckchem cat#S2742) 0 μM, 2.5 μM, 5 μM, and 10 μM; olaparib (Selleckchem cat#S1060) 0 μM, 1 μM, 5 μM, and 10 μM; mirin (Selleckchem cat#S8096) 0 μM, 25 μM, 50 μM, and 100 μM. All 0 μM cultures were treated with final 0.01% DMSO. PHA-767491(CDC7i) was added in a single time point, and cell viability and proliferation was assessed at 0, 24, 48, and 72 hrs after drug exposure. Olaparib and mirin were initially added for 48 hrs, then 250 μL of the cell suspension from each well was discarded, and 250 μL of fresh media and the appropriate drug concentrations were added back for an additional 72 hrs before cell viability counts were taken.

### Genomic DNA extraction and DNA combing

Cells were first pulse labeled for 15 minutes with 50μM CldU (Sigma), washed twice with PBS, and pulse labeled with 500μM IdU (Sigma) for 45min. Cells were centrifuged and then genomic DNA was extracted with CombHeliX DNA Extraction kit (Genomic Vision) in accordance with the manufacturer’s instructions. DNA fibers were stretched on vinyl silane-treated glass coverslips (CombiCoverslips, Genomic vision) with automated Molecular Combing System (Genomic Vision). After Combing, stretched DNA fibers were dehydrated in 37°C for 2 hours, fixed with MeOH:Acetic acid (3:1) for 10 minutes, denatured with 2.5 M HCl for 1 hour, and blocked with 5% BSA in PBST for 30 minutes. IdU and CldU were then detected with the following primary antibodies in blocking solution for 1 hour at room temperature: mouse anti-BrdU (B44) (1:25) (BD Biosciences cat#347580) for IdU and rat anti-BrdU [Bu1/75 (ICR1)] (1:25) (abcam, cat#ab6326) for CldU. After PBS wash, fibers were then incubated with secondary antibodies anti-Rat-Alexa488 (1:50) (ThermoFisher, cat#A-11006) and anti-mouse-Alexa 568 (1:50) (ThermoFisher, cat#A-11004) for 1 hour at room temperature. DNA fibers were analyzed on a Leica-DMI8 microscope at 100X and ImageJ was used to measure fiber length.

### Endpoint RT-PCR and qPCR for DYNLL1 splicing

Total RNA was isolated using RNeasy Plus Mini Kit (Qiagen), and reverse transcribed to cDNA using anchored-oligo(dT)18 primer and Transcriptor Reverse transcription (Roche). DYNLL1 was tested for splicing defects, specifically cryptic 3′ splice site usage as described(16). Primer sequences are DYNLL1-forward (5’-GTTTCGGTAGCGACGGTATCT-3’) and DYNLL1-reverse (5’-TCCGCATTTTTGATCACGGC-3’) (16). Primers were designed to obtain a PCR product that spans the spliced exon junctions in DYNLL1. The canonical splicing products results in a 133bp PCR product while the aberrant splicing product results in a 147bp PCR product. PCR reaction was carried out as follows: Initial denaturation 98°C for 30 sec, 98°C 10 sec, 62°C 30 sec, 72°C 15 sec for 40 cycles, final extension of 72°C for 10 mins. PCR products were subsequently run on a 2% agarose gel at 100V for 1.5 hours to visualize the splice isoforms. Reverse transcription–quantitative PCRs were performed and analyzed using SYBR green PCR Master Mix and a StepOnePlus Real-Time PCR system (Applied Biosystems). Primer sequences are ACTB (endogenous control) forward (5’-CACTCTTCCAGCCTTCCTTC-3’), ACTB reverse (5’-GTACAGGTCTTTGCGGATGT-3’), DYNLL1-long intron forward-(5’-ACTTGTGAGCACTCACTGAC-3’), DYNLL1-long reverse (5’-CATGGTTACCTAGTGGGCAAG-3’), DYNLL1 exon 1 forward (5’-CGCCTCAGTTTCTCTCTGTG-3’),DYNLL1 exon 1 reverse (5’-CCTGACTCCAGCTCTCCT-3’), DYNLL1 exon 2 forward (5’-AGGACATTGCGGCTCATATC-3’), DYNLL1 exon 2 reverse (5’-TTGGCCCAGGTAGAAGTAGA-3’), DYNLL1 exon 3 forward-(5’-GGGAGGAACTTCGGTAGTTATG-3’) and DYNLL1 exon 3 reverse (5’-TGGCACAGTCCATGCTTT-3’).

### RNA-Seq Analysis

RNA-Seq libraries were prepared and analyzed as described in Docking et. al (*in press*). For all MDS/AML cases in the retrospective cohort of the AML Personalized Medicine Project (AML PMP), expression of the DYNL11 RefSeq transcripts NM_001037494, NM_001037495, NM_003746 was extracted (quantified in units of transcripts per million), along with the curated SF3B1 mutation status. Differences in expression between wildtype and mutant cases were compared using two-sided *t* tests. Data for the AML PMP project is available at the European Genome-Phenome Archive under accession number EGAS00001004655.

## Supplementary Figure Legends

**Figure S1.**
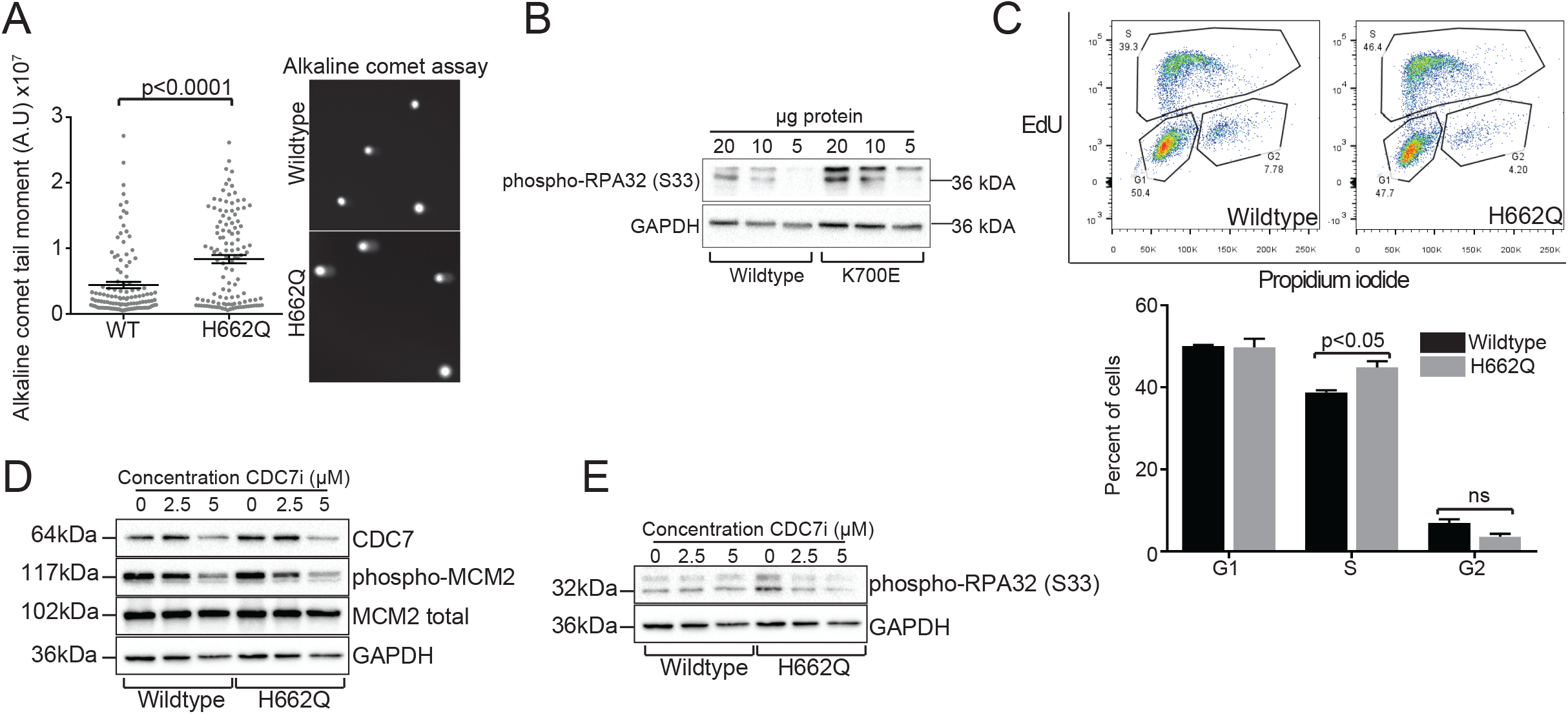
Supporting evidence and controls for replication defects in NALM6-*SF3B1*^*H662Q*^ cells (A) Alkaline comet assay for two isogenic cell line pairs with the indicated genotypes. >50 cells scored per replicate. Student’s t-test was used to calculate statistical significance. Mean values with S.E.M. error bars are shown, *n = 3*. (B) Western blot of RPA32-Ser33 phosphorylation in K562 WT and K562-*SF3B1*^*K700E*^ cells. A dilution series is shown. (C) Cell cycle progression using FACS measuring propidium iodide staining for DNA and EdU incorporation for replicating cells. Top: Representative FACs plots of wildtype (left) and SF3B1 H662Q mutant (right). Bottom: Bar graph showing quantification of FACs plots. Student’s t-test was used to calculate statistical significance. Mean values with S.E.M. error bars are shown, *n = 3*. 10,000 cells were analyzed per replicate. (D,E) Representative Western blots probing (D) CDC7, phosphorylated-MCM2, total MCM2, (E) RPA32-Ser33-P and GAPDH as a loading control in CDC7 inhibitor treated cells was used for both panels.

**Figure S2.**
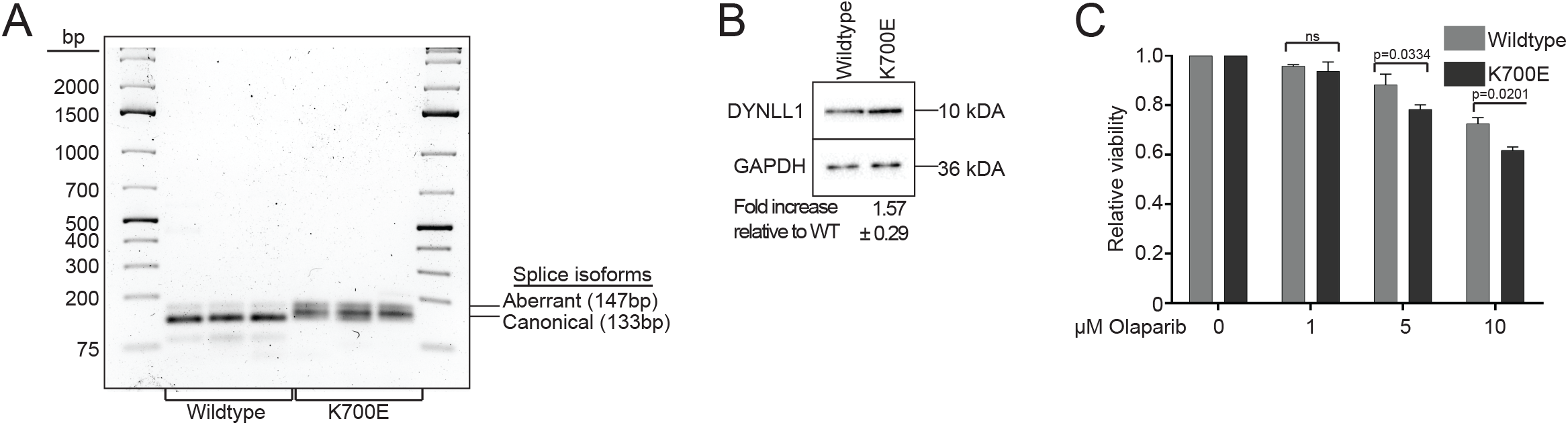
Confirmation of DYNLL1 splicing, protein, and replication stress changes in K562-*SF3B1*^*K700E*^ cells. (A) Endpoint RT-PCR analysis of DYNLL1 transcripts in the indicated cell lines. Technical triplicates are shown for each cell line. (B) Representative western blot of DYNLL1 protein, quantification of triplicates is shown below. (C) Relative cell viability of the indicated cell lines following olaparib treatment at the indicated doses. Mean values with S.E.M. error bars are shown, *n = 3*. For A-D, ANOVA with Tukey’s post-hoc test was used to determine statistical significance.

**Figure S3.**
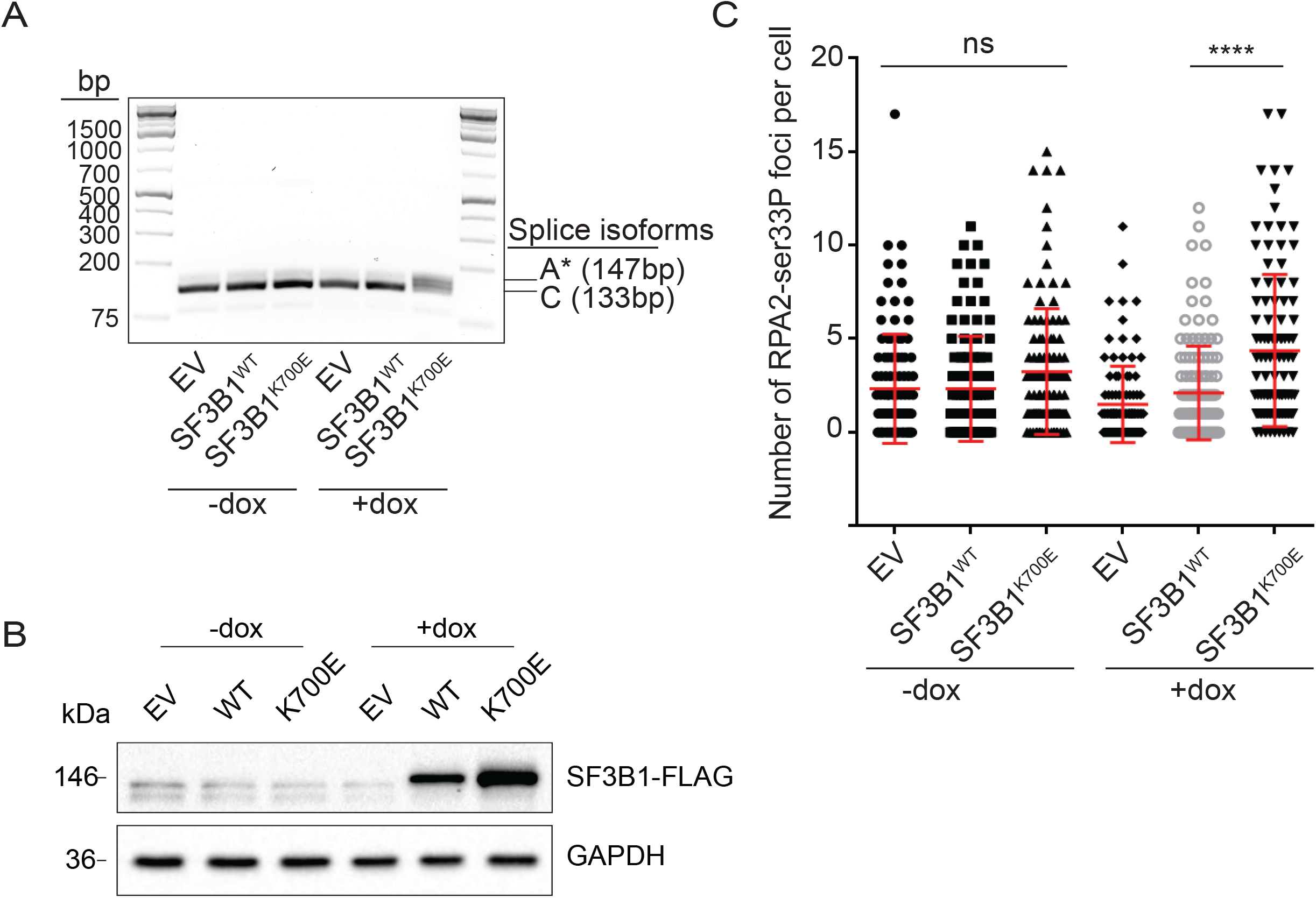
Inducible K700E model. (A) Doxycycline (dox) induction of *SF3B1*^*K700E*^ induces accumulation of the long DYNLL1 transcript. (B) SF3B1-FLAG is detectable by western blot after dox induction of the WT of K700E mutant. (C) RPA2-ser33P foci accumulation in the induced and uninduced *SF3B1*^*K700E*^ cells. Triplicate experiment. ****p<0.0001 by ANOVA. Ns = not significant.

